# A Novel Method of Immunomodulation of Endothelial cells Using Streptococcus Pyogenes and its Lysate

**DOI:** 10.1101/2020.05.13.082180

**Authors:** Mark Christopher Arokiaraj, Jarad Wilson

## Abstract

**Background:** Coronary artery diseases and autoimmune disorders are common in clinical practice. In this study, a novel method of immune-modulation to modify the endothelial function was studied to modulate the features of the endothelial cells, and thereby to reduce coronary artery disease and other disorders modulated by endothelium.

**Methods:** HUVEC cells were seeded in the cell culture, and streptococcus pyogenes were added to the cell culture, and the supernatant was studied for the secreted proteins. In the second phase, the bacterial lysate was synthesized, and the lysate was added to cell culture; and the proteins in the supernatant were studied at various time intervals.

**Results:** When streptococcus pyogenes alone was added to culture, E Cadherin, Angiostatin, EpCAM and PDGF-AB were some of the biomarkers elevated significantly. HCC1, IGFBP2 and TIMP were some of the biomarkers which showed a reduction. When the lysate was added, the cell-culture was maintained for a longer time, and it showed the synthesis of immune regulatory cytokines. Heatmap analysis showed a significant number of proteins/cytokines concerning the immune/pathways, and toll-like receptors superfamily were modified. BLC, IL 17, BMP 7, PARC, Contactin2, IL 10 Rb, NAP 2 (CXCL 7), Eotaxin 2 were maximally increased. By principal component analysis, the results observed were significant.

**Conclusion:** There is potential for a novel method of immunomodulation of the endothelial cells, which have pleiotropic functions, using streptococcus pyogenes and its lysates.

## Background

Immune-related disorders are common in clinical practice. Though commonly handled disease by the immune system is an infection, most of the other disorders like autoimmune disorders, leukemias, and even non-communicable diseases have an immune basis. A broad-spectrum treatment method to modulate immune disorders would be useful.

Rheumatic heart diseases are common in the general population in the Asian countries with a prevalence of about 7.7 to 51/1000 community based on echocardiography screening in school in India.^1^ Streptococcus pyogenes infections are associated with sore-throats, and they are also associated with rheumatic heart diseases. Rheumatic fever is associated with migratory joint pains and occasionally pan-carditis. In patients with rheumatic heart diseases, the incidence of coronary artery diseases is low.^2-5^ This was observed in some studies in India and neighboring countries. Hence, in this study the streptococcus pyogenes infections were used to study the immune modulatory potentials in the endothelial cells.

Streptococcus pyogenes are known to produce a unique enzyme that is useful in cleaving immunoglobulin G in the blood (Ides).^6-8^ Its usefulness therapeutically has been tested to cleave IgG antibodies.^9,10^ Streptococcus pyogenes secrete serum optical factor, which shows increased uptake of HDL, and thereby it can reduce atherosclerosis.^11,12^

In this study, streptococcus pyogenes was used to infect the endothelial cells, and the endothelial response was evaluated. Also, in the later part of the study the streptococcus pyogenes’ lysate was used on the endothelial cells, and the results were evaluated. Microorganisms and their diseases and susceptibility to infections can prevent autoimmune disorders though this phenomenon is not well studied.^13^ The incidence of autoimmune disorders is common in the migrant populations especially of Asian ethnicity.^14,15^ This is commonly attributed as the hygiene hypothesis, and the exact mechanism is speculative and not decisively studied.

In our center out of 650 consecutive valve replacement surgeries six cases underwent concomitant coronary artery bypass surgery. Out of these 6 cases a clear etiology of rheumatic heart disease was not established on any of the cases, and the primary etiologies were ischemia MR and degenerative sclerotic calcific aortic valve. The study was performed in search of novel applications of streptococcus pyogenes in regulating immune functions, and its related effects on cardiovascular and its pleiotropic functions.

## Methods

Streptococcus pyogenes were obtained from the laboratory (Streptococcus pyogenes Rosenbach – ATCC,19615TM Lancefield Group A). The bacteria were cultured by seeding, and the colonies were inoculated with endothelial cells. Serial observations about the endothelial cells were made at regular intervals by microscopy. The supernatant was collected, and inflammatory markers were studied at serial time intervals.

In the second phase of the study, the streptococcus pyogenes’ lysate was prepared from the same bacteria. The lysate was added to the endothelial cells, and the endothelial cell response was studied at serial intervals -0, 36,138, 336, 672 hours, and control samples were studied. The biomarkers secreted were studied, and the results were achieved.

### Statistical Analysis - Data filtration

The biomarkers demonstrating no variation among all the samples (zero variance) were excluded from the data profile analysis since they do not contribute regarding distinguishing samples from each other.

### Heatmap

The biomarker values were standardized (centering and scaling) by subtracting the average and then dividing by the standard deviation. The standardized data were plotted in a heatmap with hierarchical clustering by Euclidean distance.

### Principal Component analysis

The various expression levels of multiple biomarkers may come from a common underlying factor/mechanism. The principal component analysis (PCA) decompose the data set into different principal components (PCs) sorted by their contribution to total variance/variation in the dataset. These PCs are linear transformations/combinations of standardized biomarker values. By observing the location of a sample on the plot of the first 2 PCs explaining the most variation, we can tell the pattern of samples.

### Software

All the analyses were conducted in the R programming language V3.6.0 (R Core Team 2017).

## Results

### Direct Streptococcus pyogenes’ effects

Tables 1 to 5 summarize the effects of the bacteria on the cell culture of endothelial cells. There is a significant reduction in HCC1, IGFBP2, PDGFAA, and TIMP. Macrophage inhibiting factor and lymphocyte inhibition factors showed a decrease in the levels. There is a marked increase in E Cadherin, Angiostatin, DAN, Ep-CAM, CFG RIIBC, PDGF-AB, gp 130, TPO, Tie 2, and Angiogenin. ICAM-1, IL6, Endoglin, Trail 3, and PRECAM-1 showed an increasing trend.

**Table 1.**

| LOD | MAX | (pg/ml) | 0 hour treated | 2 hour treated | 6 hour treated | 10 hour treated | 14 hour treated | (pg/ml) | 14 hour untreated |
| --- | --- | --- | --- | --- | --- | --- | --- | --- | --- |
| 14.2 | 40,000.0 | 6Ckine | 1.9 | 1.4 | 6.2 | 0.3 | 0.0 | 6Ckine | 2.0 |
| 10.9 | 4,000.0 | Axl | 0.0 | 0.0 | 0.0 | 16.1 | 14.8 | Axl | 11.8 |
| 17.8 | 20,000.0 | BTC | 0.0 | 0.0 | 0.0 | 0.0 | 0.0 | BTC | 0.0 |
| 26.2 | 40,000.0 | CCL28 | 0.0 | 0.0 | 0.0 | 0.0 | 0.0 | CCL28 | 0.0 |
| 26.7 | 50,000.0 | CTACK | 0.0 | 0.0 | 0.0 | 0.0 | 0.0 | CTACK | 0.0 |
| 11.7 | 20,000.0 | CXCL16 | 0.0 | 6.6 | 0.4 | 6.4 | 19.2 | CXCL16 | 36.4 |
| 13.7 | 10,000.0 | ENA-78 | 2.1 | 0.0 | 0.0 | 0.0 | 9.0 | ENA-78 | 2.0 |
| 62.4 | 20,000.0 | Eotaxin-3 | 0.0 | 0.0 | 0.0 | 0.0 | 0.0 | Eotaxin-3 | 0.0 |
| 4.2 | 10,000.0 | GCP-2 | 0.0 | 0.0 | 0.0 | 0.0 | 0.0 | GCP-2 | 2.8 |
| 2.9 | 1,000.0 | GRO | 1.4 | 3.2 | 4.1 | 6.9 | 8.4 | GRO | 25.0 |
| 6.7 | 4,000.0 | HCC-1 | 0.0 | 7.4 | 44.9 | 69.6 | 90.9 | HCC-1 | 213.1 |
| 4.0 | 3,333.3 | HCC-4 | 0.0 | 0.1 | 0.0 | 0.4 | 0.4 | HCC-4 | 0.2 |
| 800.0 | 200,000.0 | IL-9 | 0.0 | 206.2 | 28.9 | 0.0 | 109.6 | IL-9 | 0.0 |
| 49.6 | 100,000.0 | IL-17F | 0.1 | 0.1 | 0.3 | 0.1 | 0.4 | IL-17F | 0.0 |
| 52.0 | 60,000.0 | IL-18 BPα | 4.0 | 3.4 | 0.0 | 0.0 | 33.3 | IL-18 BPα | 0.0 |
| 7.7 | 10,000.0 | IL-28A | 0.0 | 0.0 | 0.5 | 0.0 | 0.0 | IL-28A | 0.0 |
| 302.0 | 100,000.0 | IL-29 | 0.0 | 30.5 | 0.0 | 0.0 | 0.0 | IL-29 | 0.0 |
| 97.8 | 40,000.0 | IL-31 | 0.0 | 0.0 | 0.0 | 0.0 | 7.8 | IL-31 | 0.0 |
| 13.7 | 10,000.0 | IP-10 | 0.5 | 0.0 | 1.8 | 0.0 | 0.9 | IP-10 | 0.1 |
| 16.8 | 10,000.0 | I-TAC | 0.0 | 2.4 | 0.0 | 0.0 | 5.9 | I-TAC | 9.7 |
| 28.4 | 13,000.0 | LIF | 0.0 | 16.4 | 18.8 | 0.2 | 47.9 | LIF | 0.0 |
| 15.6 | 10,000.0 | LIGHT | 0.0 | 0.0 | 0.0 | 0.0 | 0.9 | LIGHT | 0.0 |
| 47.5 | 100,000.0 | Lymphotactin | 31.3 | 43.7 | 45.7 | 0.0 | 78.8 | Lymphotactin | 49.3 |
| 5.7 | 2,000.0 | MCP-2 | 0.9 | 0.0 | 0.7 | 0.0 | 1.7 | MCP-2 | 1.7 |
| 4.2 | 4,000.0 | MCP-3 | 0.0 | 0.0 | 0.0 | 0.0 | 0.0 | MCP-3 | 0.0 |
| 4.6 | 1,111.1 | MCP-4 | 0.0 | 0.1 | 0.4 | 0.0 | 0.1 | MCP-4 | 0.5 |
| 4.9 | 1,111.1 | MDC | 0.0 | 0.0 | 0.0 | 0.0 | 0.3 | MDC | 0.2 |
| 13.6 | 4,000.0 | MIF | 72.4 | 70.9 | 323.2 | 709.2 | 1,417.9 | MIF | 74.6 |
| 4.5 | 4,000.0 | MIP-3a | 0.0 | 0.5 | 0.7 | 0.5 | 2.5 | MIP-3a | 0.8 |
| 17.8 | 20,000.0 | MIP-3b | 0.0 | 0.0 | 0.0 | 0.0 | 1.6 | MIP-3b | 0.0 |
| 15.0 | 10,000.0 | MPIF-1 | 0.0 | 0.0 | 1.7 | 2.0 | 0.1 | MPIF-1 | 0.7 |
| 213.2 | 100,000.0 | MSP | 0.0 | 2.4 | 39.5 | 17.7 | 61.1 | MSP | 21.1 |
| 1.9 | 444.4 | NAP-2 | 0.0 | 0.0 | 0.3 | 0.0 | 0.1 | NAP-2 | 0.1 |
| 28.6 | 100,000.0 | OPN | 37.1 | 52.9 | 36.9 | 1.8 | 36.8 | OPN | 14.2 |
| 4.9 | 4,000.0 | PARC | 0.0 | 0.2 | 0.4 | 0.6 | 0.5 | PARC | 1.2 |
| 219.7 | 100,000.0 | PF4 | 0.0 | 0.0 | 0.0 | 0.0 | 0.0 | PF4 | 0.4 |
| 2.8 | 10,000.0 | SDF-1α | 0.1 | 0.2 | 0.3 | 0.1 | 2.8 | SDF-1α | 1.4 |
| 6.9 | 10,000.0 | TARC | 0.0 | 0.0 | 0.0 | 0.0 | 0.0 | TARC | 0.9 |
| 85.2 | 100,000.0 | TECK | 0.0 | 0.0 | 0.0 | 0.0 | 38.0 | TECK | 0.0 |
| 7.3 | 10,000.0 | TSLP | 0.0 | 0.0 | 0.4 | 0.0 | 2.9 | TSLP | 10.2 |

**Table 2.**
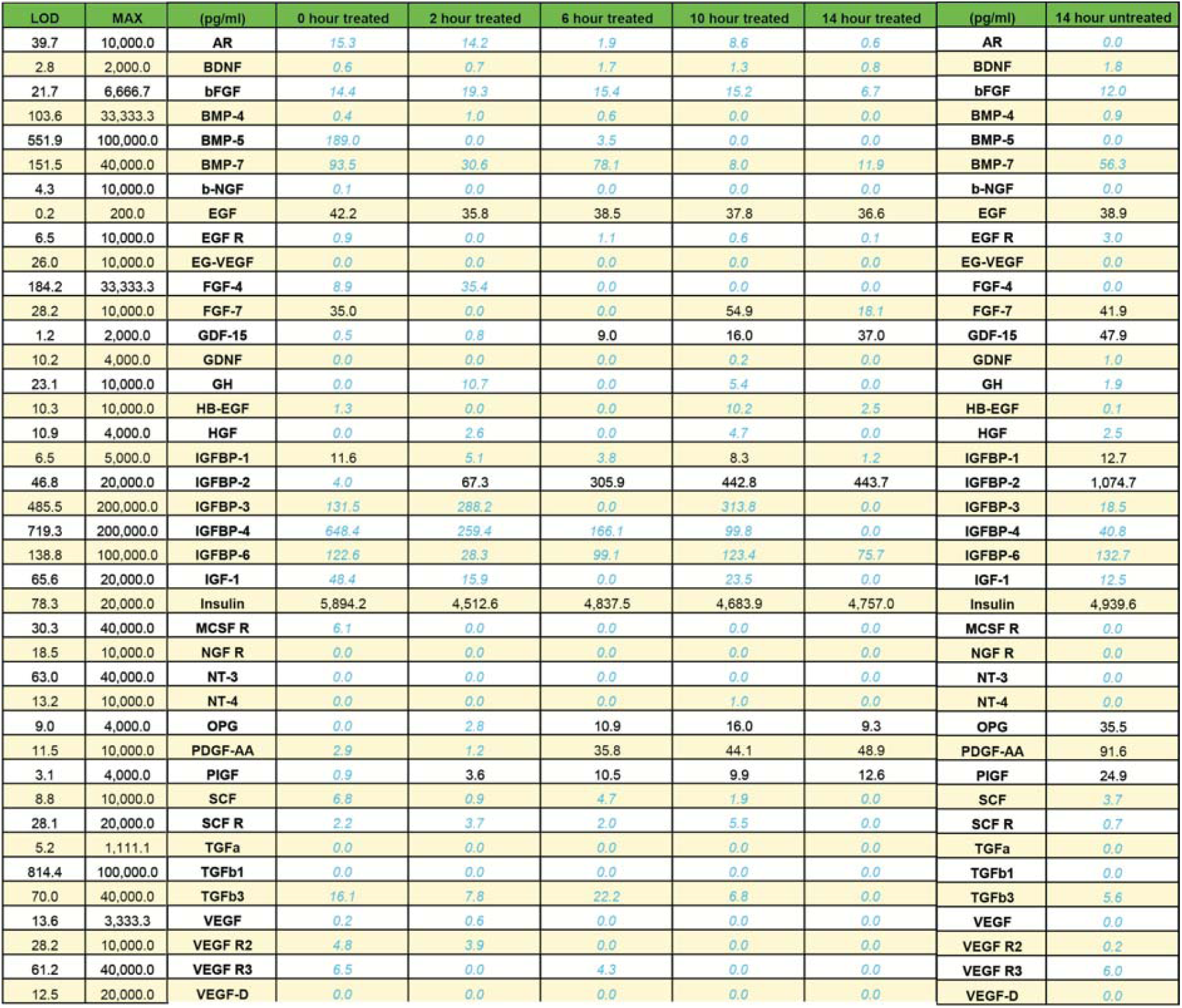

**Table 3.**

| LOD | MAX | (pg/ml) | 0 hour treated | 2 hour treated | 6 hour treated | 10 hour treated | 14 hour treated | (pg/ml) | 14 hour untreated |
| --- | --- | --- | --- | --- | --- | --- | --- | --- | --- |
| 263.8 | 11,111.1 | Activin A | 346.3 | 251.0 | 343.9 | 281.7 | 472.9 | Activin A | 209.1 |
| 16.8 | 10,000.0 | AgRP | 30.3 | 16.5 | 45.4 | 18.9 | 184.3 | AgRP | 20.0 |
| 5.5 | 2,000.0 | Angiogenin | 37.9 | 29.9 | 43.4 | 29.8 | 144.1 | Angiogenin | 26.2 |
| 56.3 | 40,000.0 | ANG-1 | 246.5 | 209.1 | 257.7 | 258.2 | 322.4 | ANG-1 | 163.3 |
| 2,267.7 | 1,000,000.0 | Angiostatin | 4,942.9 | 3,576.9 | 8,953.9 | 3,369.3 | 40,517.2 | Angiostatin | 3,978.9 |
| 10.4 | 10,000.0 | Cathepsin S | 25.7 | 19.2 | 27.9 | 26.0 | 184.6 | Cathepsin S | 17.8 |
| 10.4 | 10,000.0 | CD40 | 11.0 | 13.1 | 23.3 | 23.6 | 37.7 | CD40 | 27.8 |
| 23.1 | 10,000.0 | Cripto-1 | 5.2 | 0.0 | 20.4 | 0.0 | 35.2 | Cripto-1 | 0.0 |
| 169.4 | 40,000.0 | DAN | 374.8 | 70.5 | 562.2 | 165.8 | 11,816.4 | DAN | 119.4 |
| 49.8 | 8,888.9 | DKK-1 | 35.7 | 192.3 | 1,122.9 | 1,466.7 | 2,365.2 | DKK-1 | 3,009.6 |
| 173.9 | 80,000.0 | E-Cadherin | 333.6 | 149.1 | 1,478.0 | 128.0 | 4,275.4 | E-Cadherin | 167.7 |
| 13.7 | 20,000.0 | EpCAM | 8.0 | 0.5 | 17.5 | 4.1 | 189.7 | EpCAM | 2.8 |
| 6.3 | 2,000.0 | FAS L | 20.3 | 4.6 | 19.0 | 5.0 | 40.8 | FAS L | 12.1 |
| 58.0 | 10,000.0 | Fcg RIIBC | 128.9 | 43.3 | 323.8 | 60.5 | 568.9 | Fcg RIIBC | 134.4 |
| 22.8 | 40,000.0 | Follistatin | 2.7 | 5.5 | 15.2 | 23.0 | 86.0 | Follistatin | 63.2 |
| 102.6 | 100,000.0 | Galectin-7 | 25.0 | 48.2 | 58.6 | 21.9 | 56.5 | Galectin-7 | 81.9 |
| 235.1 | 100,000.0 | ICAM-2 | 433.2 | 203.9 | 3,766.3 | 3,949.3 | 9,610.2 | ICAM-2 | 2,397.8 |
| 50.2 | 10,000.0 | IL-13 R1 | 534.1 | 531.2 | 1,004.0 | 567.2 | 2,667.3 | IL-13 R1 | 444.9 |
| 70.7 | 20,000.0 | IL-13 R2 | 140.1 | 105.9 | 153.7 | 228.2 | 155.1 | IL-13 R2 | 119.3 |
| 103.7 | 40,000.0 | IL-17B | 310.7 | 241.8 | 1,552.4 | 384.4 | 551.4 | IL-17B | 179.7 |
| 22.3 | 10,000.0 | IL-2 Ra | 98.5 | 57.8 | 108.3 | 88.1 | 207.3 | IL-2 Ra | 19.1 |
| 253.8 | 100,000.0 | IL-2 Rb | 199.8 | 134.6 | 170.4 | 265.7 | 191.3 | IL-2 Rb | 206.8 |
| 171.6 | 40,000.0 | IL-23 | 0.0 | 4.7 | 122.3 | 26.1 | 59.0 | IL-23 | 0.0 |
| 8.2 | 4,000.0 | LAP(TGFb1) | 34.5 | 66.5 | 188.4 | 175.7 | 267.7 | LAP(TGFb1) | 211.7 |
| 48.7 | 20,000.0 | NrCAM | 112.7 | 47.1 | 125.4 | 91.0 | 71.7 | NrCAM | 55.0 |
| 107.8 | 40,000.0 | PAI-1 | 1,907.2 | 17,224.8 | 23,895.7 | 22,084.3 | 20,426.1 | PAI-1 | 16,711.3 |
| 28.4 | 10,000.0 | PDGF-AB | 45.6 | 28.4 | 92.9 | 63.5 | 138.1 | PDGF-AB | 15.5 |
| 70.9 | 20,000.0 | Resistin | 49.0 | 4.1 | 56.6 | 26.7 | 49.8 | Resistin | 9.2 |
| 32.7 | 13,333.3 | SDF-1b | 14.9 | 6.3 | 52.0 | 5.0 | 10.3 | SDF-1b | 1.3 |
| 239.7 | 80,000.0 | gp130 | 330.2 | 37.4 | 859.8 | 320.1 | 892.3 | gp130 | 16.3 |
| 25.2 | 40,000.0 | Shh-N | 11.6 | 14.8 | 31.2 | 32.8 | 42.2 | Shh-N | 54.0 |
| 32.2 | 10,000.0 | Siglec-5 | 91.6 | 10.7 | 344.8 | 45.1 | 95.8 | Siglec-5 | 38.5 |
| 14.2 | 4,000.0 | ST2 | 28.6 | 0.9 | 27.7 | 17.1 | 31.1 | ST2 | 35.0 |
| 104.8 | 40,000.0 | TGFb2 | 57.1 | 18.1 | 85.6 | 54.4 | 72.2 | TGFb2 | 42.9 |
| 46.5 | 10,000.0 | Tie-2 | 122.7 | 22.5 | 451.4 | 51.2 | 79.3 | Tie-2 | 27.0 |
| 335.0 | 200,000.0 | TPO | 249.1 | 66.3 | 155.7 | 51.1 | 212.2 | TPO | 57.5 |
| 26.3 | 8,000.0 | TRAIL R4 | 61.5 | 38.7 | 54.8 | 33.1 | 58.3 | TRAIL R4 | 31.5 |
| 111.9 | 20,000.0 | TREM-1 | 468.6 | 178.7 | 2,561.2 | 209.8 | 468.1 | TREM-1 | 167.4 |
| 26.4 | 20,000.0 | VEGF-C | 57.8 | 1.9 | 37.9 | 35.5 | 37.3 | VEGF-C | 19.6 |
| 118.8 | 40,000.0 | VEGF R1 | 106.8 | 593.6 | 2,745.2 | 3,514.9 | 4,425.7 | VEGF R1 | 8,579.2 |

**Table 4.**

| LOD | MAX | (pg/ml) | 0 hour treated | 2 hour treated | 6 hour treated | 10 hour treated | 14 hour treated | (pg/ml) | 14 hour untreated |
| --- | --- | --- | --- | --- | --- | --- | --- | --- | --- |
| 2.3 | 666.7 | BLC | 0.0 | 0.0 | 0.0 | 0.0 | 0.0 | BLC | 0.0 |
| 15.4 | 4,000.0 | Eotaxin | 0.0 | 0.0 | 0.0 | 0.0 | 0.0 | Eotaxin | 0.0 |
| 11.9 | 1,000.0 | Eotaxin-2 | 0.0 | 0.0 | 0.0 | 0.0 | 0.3 | Eotaxin-2 | 0.0 |
| 35.0 | 20,000.0 | G-CSF | 0.0 | 0.0 | 0.0 | 0.0 | 0.1 | G-CSF | 2.0 |
| 4.8 | 1,000.0 | GM-CSF | 0.0 | 0.0 | 0.0 | 3.4 | 6.1 | GM-CSF | 0.9 |
| 14.8 | 4,000.0 | I-309 | 0.0 | 0.0 | 0.0 | 1.1 | 1.6 | I-309 | 0.0 |
| 56.5 | 33,333.3 | ICAM-1 | 23.2 | 54.7 | 69.6 | 76.2 | 159.3 | ICAM-1 | 52.0 |
| 14.2 | 2,000.0 | IFNγ | 0.0 | 0.0 | 0.0 | 0.0 | 7.0 | IFNγ | 0.0 |
| 5.4 | 2,000.0 | IL-1a | 0.0 | 0.0 | 2.8 | 6.0 | 15.0 | IL-1a | 11.4 |
| 2.3 | 1,000.0 | IL-1b | 0.0 | 0.0 | 0.0 | 0.0 | 0.0 | IL-1b | 0.0 |
| 5.7 | 222.2 | IL-1ra | 0.0 | 0.0 | 0.0 | 0.0 | 0.0 | IL-1ra | 7.2 |
| 6.8 | 2,000.0 | IL-2 | 0.8 | 0.0 | 0.0 | 0.0 | 0.0 | IL-2 | 4.5 |
| 5.2 | 2,000.0 | IL-4 | 0.0 | 0.0 | 0.0 | 0.0 | 0.0 | IL-4 | 0.0 |
| 17.3 | 4,000.0 | IL-5 | 0.0 | 2.7 | 0.0 | 0.0 | 0.0 | IL-5 | 8.7 |
| 8.1 | 2,000.0 | IL-6 | 0.0 | 9.2 | 22.7 | 77.4 | 136.3 | IL-6 | 53.8 |
| 19.5 | 10,000.0 | IL-6R | 0.0 | 0.0 | 0.0 | 0.0 | 0.0 | IL-6R | 0.0 |
| 14.0 | 4,000.0 | IL-7 | 0.0 | 3.3 | 0.0 | 0.0 | 0.0 | IL-7 | 9.8 |
| 2.1 | 500.0 | IL-8 | 4.6 | 16.5 | 7.0 | 4.9 | 17.2 | IL-8 | 128.7 |
| 8.8 | 4,000.0 | IL-10 | 0.0 | 0.8 | 0.0 | 0.0 | 0.1 | IL-10 | 2.4 |
| 44.0 | 20,000.0 | IL-11 | 0.0 | 0.0 | 0.0 | 0.0 | 3.8 | IL-11 | 0.0 |
| 17.9 | 10,000.0 | IL-12p40 | 0.0 | 0.0 | 0.0 | 0.0 | 0.0 | IL-12p40 | 0.0 |
| 1.2 | 500.0 | IL-12p70 | 0.3 | 0.5 | 0.2 | 0.0 | 1.0 | IL-12p70 | 0.6 |
| 17.2 | 1,000.0 | IL-13 | 4.2 | 9.1 | 23.0 | 23.5 | 69.3 | IL-13 | 97.3 |
| 41.0 | 4,000.0 | IL-15 | 0.0 | 0.0 | 0.0 | 4.5 | 27.9 | IL-15 | 32.1 |
| 14.8 | 5,000.0 | IL-16 | 0.0 | 0.0 | 0.0 | 0.0 | 0.0 | IL-16 | 1.0 |
| 11.1 | 4,000.0 | IL-17 | 0.0 | 0.0 | 0.0 | 0.0 | 0.6 | IL-17 | 0.0 |
| 15.2 | 2,000.0 | MCP-1 | 7.2 | 24.0 | 100.7 | 276.7 | 266.4 | MCP-1 | 219.0 |
| 3.0 | 4,000.0 | MCSF | 0.0 | 0.0 | 0.0 | 0.0 | 0.0 | MCSF | 0.0 |
| 11.2 | 5,000.0 | MIG | 0.0 | 0.0 | 0.0 | 0.0 | 0.6 | MIG | 0.2 |
| 17.1 | 3,333.3 | MIP-1a | 0.0 | 0.0 | 0.0 | 0.0 | 0.0 | MIP-1a | 0.0 |
| 2.8 | 1,000.0 | MIP-1b | 0.0 | 0.0 | 0.0 | 0.0 | 0.0 | MIP-1b | 0.0 |
| 6.7 | 3,333.3 | MIP-1d | 0.0 | 0.0 | 0.0 | 0.0 | 0.0 | MIP-1d | 0.0 |
| 4.0 | 2,000.0 | PDGF-BB | 0.0 | 0.0 | 0.9 | 3.9 | 9.6 | PDGF-BB | 5.7 |
| 29.9 | 6,666.7 | RANTES | 0.0 | 0.0 | 0.0 | 0.0 | 0.0 | RANTES | 0.0 |
| 89.8 | 13,333.3 | TIMP-1 | 0.0 | 1,286.7 | 2,866.2 | 2,781.3 | 3,291.9 | TIMP-1 | 3,752.9 |
| 24.5 | 40,000.0 | TIMP-2 | 0.0 | 53.1 | 359.7 | 444.6 | 566.4 | TIMP-2 | 1,439.1 |
| 37.2 | 2,000.0 | TNFa | 0.0 | 44.1 | 18.3 | 49.4 | 41.8 | TNFa | 55.4 |
| 66.1 | 20,000.0 | TNFb | 0.0 | 6.2 | 0.0 | 13.5 | 19.9 | TNFb | 2.3 |
| 36.6 | 40,000.0 | TNF RI | 0.0 | 0.0 | 7.5 | 1.2 | 10.8 | TNF RI | 7.0 |
| 118.3 | 40,000.0 | TNF RII | 0.0 | 0.0 | 0.0 | 0.0 | 9.1 | TNF RII | 0.0 |

**Table 5.**
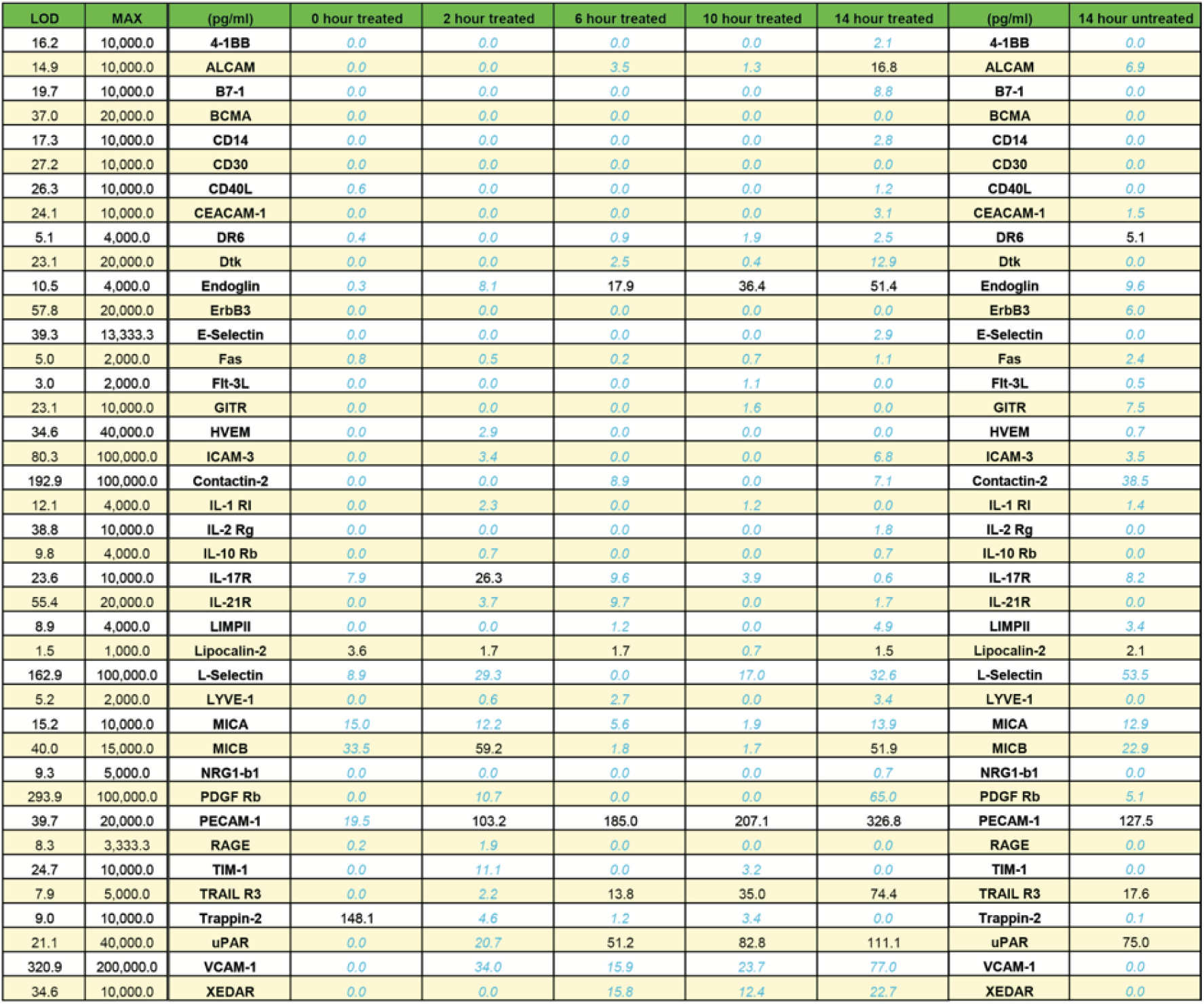

### Streptococcus pyogenes’ Lysate Challenge

The results of the streptococcal lysate challenge on the endothelial cells are summarized in the heat map results (Figure 1). BLC, IL 17, BMP 7, PARC, Contactin2, IL 10 Rb, NAP 2 (CXCL 7), Eotaxin 2 were maximally increased. Other secreted markers that were induced – trappin2, SDF-1a, FGF7, CCL 28, 4-1BB, VCAM 1, VEGF R3, GITR, SIGLEC 5, IL13 R1, CD30, TGF B2 and GP 130. The markers which were unequivocally reduced were MIF, Fas, IL13R2, TIMP2, Follistatin, IGFBP-2, DKK1, TNF R2, Hb EGF. There are 18 biomarkers showing zero variance among all the samples, including BTC, IL-9, IL-29, MCP-4, CD40, DAN, E-Cadherin, b-NGF, EGF R, EG-VEGF, FGF-4, IGF-1, NT-3, SCF, TGFb3, IL-5, BCMA, and E-Selectin. These biomarkers were excluded from the analysis. Details of the principal components and the results are shown in figure 1 and 2 (and supplement file). The results of the principal component analysis are significant.

**Figure 1A.**
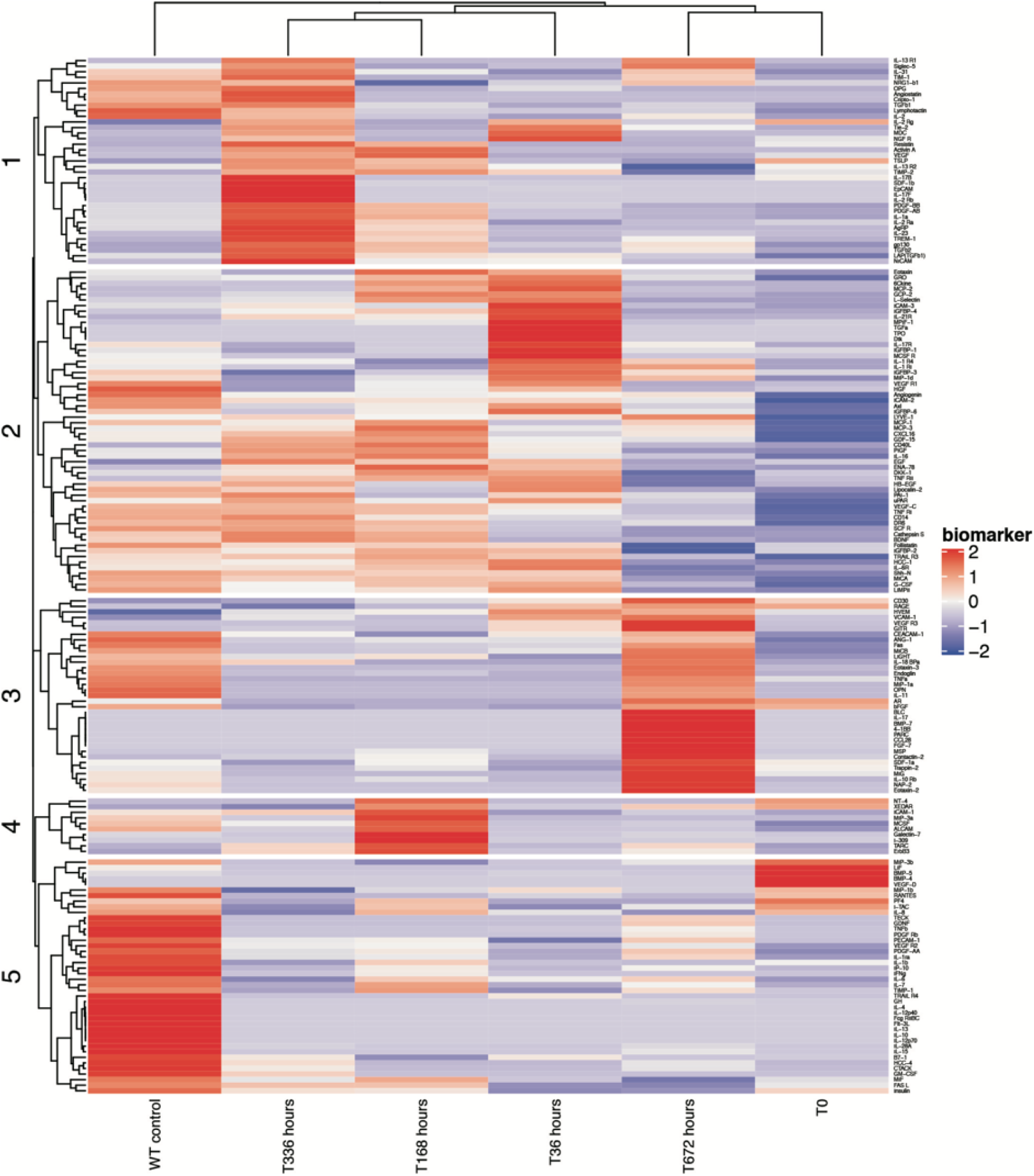
Heat map and the classification of the principal components (1-5) for analysis of the biomarkers after streptococcus pyogenes lysate treatment on the endothelial cells.

**Figure 1B.**
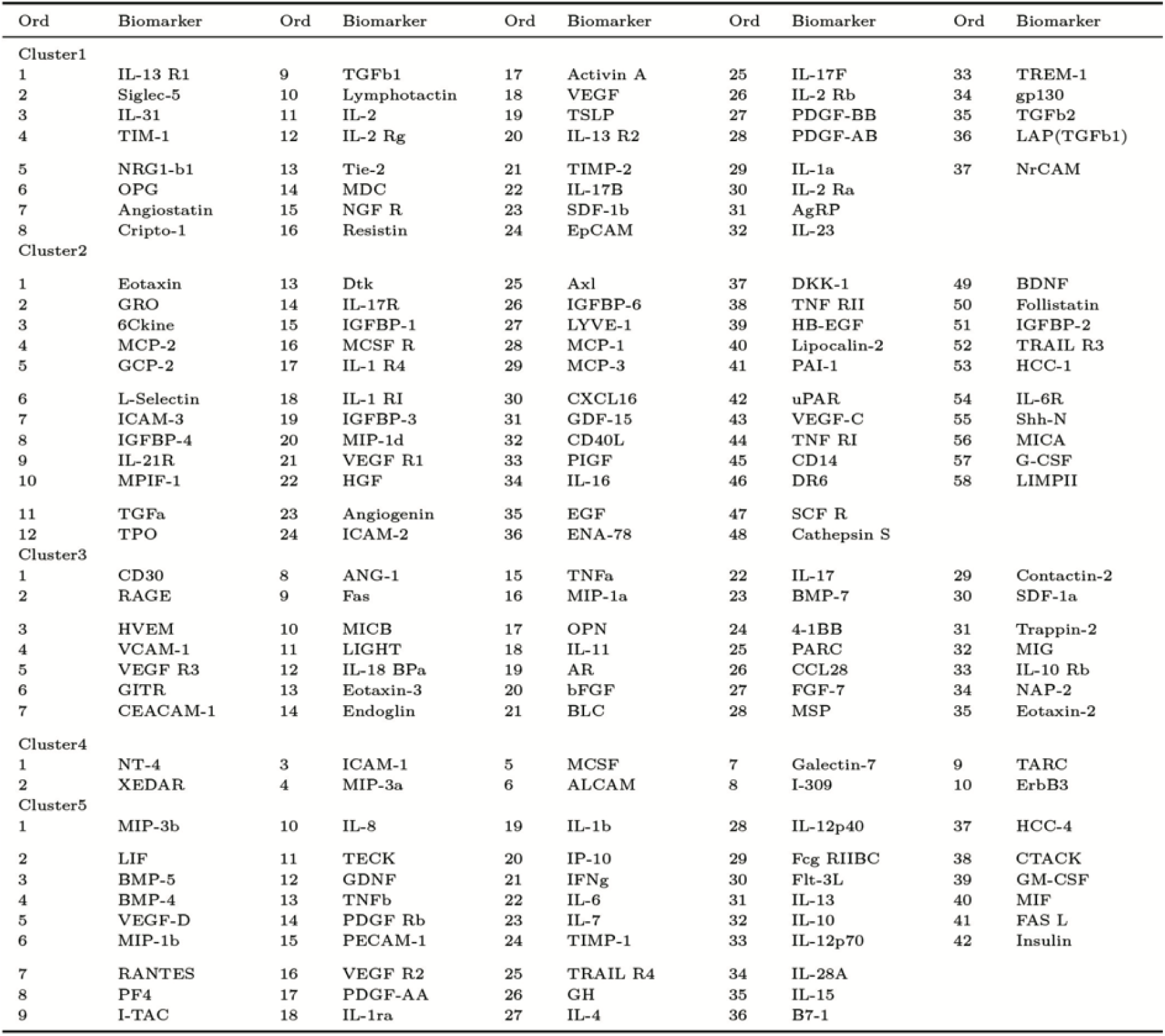
Legends for Figure 1.

**Figure 2:**
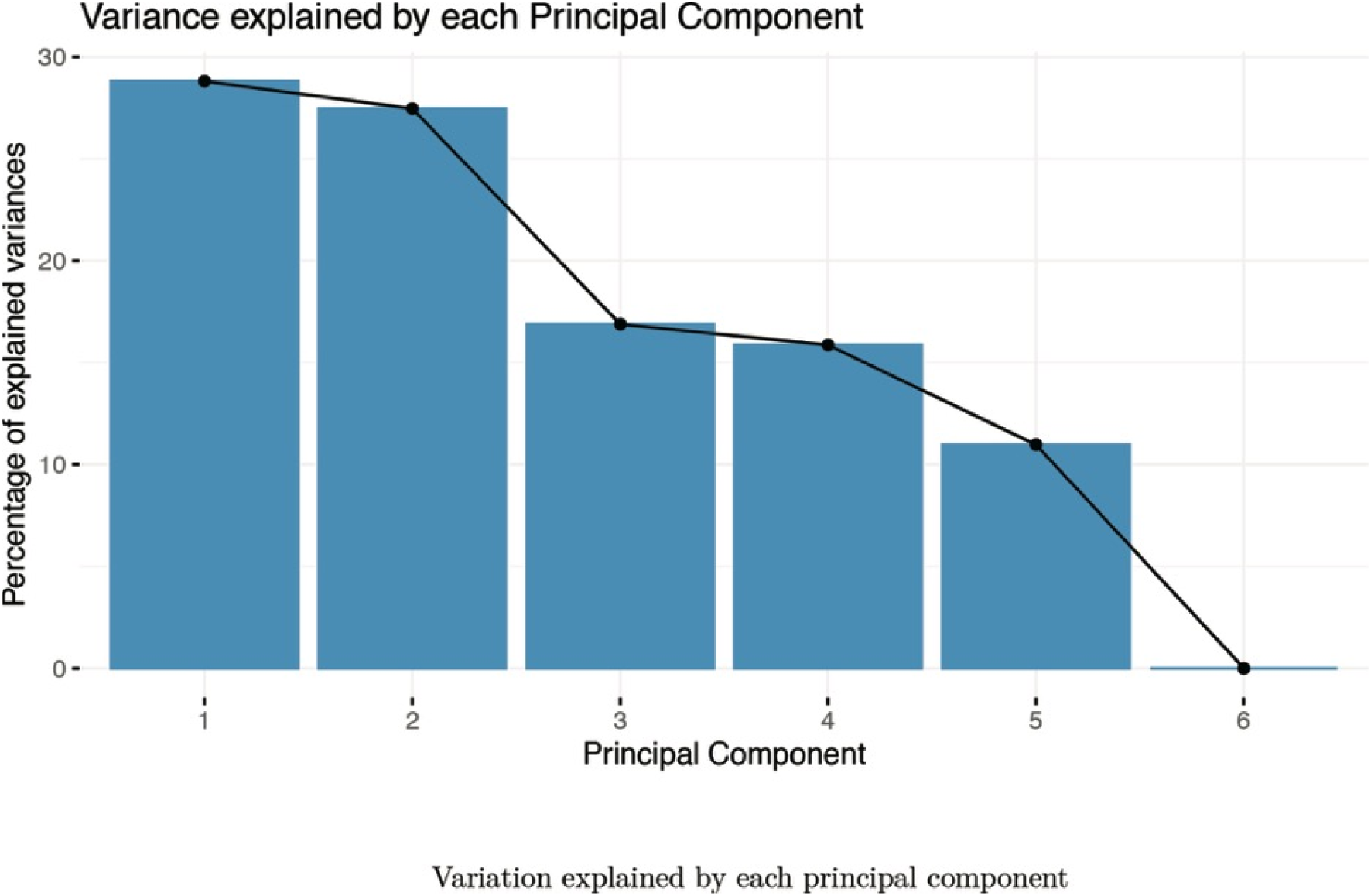
Shows the variance explained by each principal component.

## Discussion

The incidence of rheumatic heart disease is showing a decreasing trend, and the incidence of coronary artery disease is rising in the recent days. The incidence in our center and other centers as well show a low association of coronary artery disease with rheumatic valvular heart diseases, irrespective of age and metabolic characteristics.^2,3^ The incidence of mixed procedure i.e., combined valve surgeries and bypass surgeries, is <5%. And further analysis looking for rheumatic valvular etiology in these combined procedures would be much lesser, even in high volume centers like TUMS.^4^

### Direct streptococcus pyogenes immune response - Decreased biomarkers

Specific biomarkers like HCC1, IGFBP2, PDGF-AA, and TIMP decrease in the levels compared to controls. Hemofiltrate CC is a chemokine, which attracts and acts through CCR1 receptors.^16^ It is widely secreted by various tissues. Insulin-like growth factor binding protein 2 reduces the risk of diabetes. IGFBP2 is implicated in the regulation of IGF in most tissues.^17^ Blocking of IGF BP2 results in the reduction of tumor and metastasis.^18^ Platelet-derived growth factor AA is involved in the migration of smooth muscle cells.^19-21^ Reduction in the TIMP metallopeptidase inhibitor1 and also has anti-angiogenic activity is associated with a reduction in the adverse clinical events in acute kidney injuries.^22^ MIF (macrophage migration inhibitory factor) and LIF (leukemia inhibitory factor) levels were reduced. MIF is a widely expressed pleiotropic cytokine, and it is involved in the stimulation of other inflammatory cytokines like TNF alpha, INF gamma, IL6, IL 12, CXCL8 etc.^23^ LIF is involved in cell differentiation, and maturation and stimulation lead to JAK/STAT and MAPK cascades.^24^

### Increased biomarkers with direct bacterial infection

Increased activity was found in E Cadherin is involved in cell-to-cell interactions, and they have tumor suppressor effects.^25^ Angiostatin, DAN, is an inhibitor of BMP and TNF. Angiostatin is engaged in the reduction of angiogenesis.^26,27^ There is a marked increase in EPCAM, which are complex proteins that promote transcription factor-mediated pluripotency reprogramming,^28^ CFG RIIC and PDGF AB.^29^ The Platelet-derived growth factor has active angiogenic potential and mitogenesis and acts on various tissues. Angiogenin may maintain blood homeostasis and participates in anti-inflammatory activity and has antibacterial and antiviral properties.^30^

### Streptococcus pyogenes’ lysate effects and Heat map analysis

When the cells were treated with streptococcus pyogenes’ lysate, the levels of BLC -the B lymphocyte chemoattractant protein (CXCL13) was increased.^31^ Contactin 2 is a neuronal membrane protein, and it acts as an active cell adhesion molecule.^32-34^ IL 17 is an inflammatory protein, and it was induced after lysate treatment. IL17 induces the production of GCSF and chemokines like CXCL1 and 2.^35^ IL17 is strongly associated with chronic inflammation associated with autoimmune disorders. PARC (parkin like ubiquitin ligase) is a cytoplasmic anchor protein to p53-associated protein complexes.^36-38^

CXCL7 is involved in neutrophil chemotaxis, adhesion to the endothelial cells, and trans-endothelial migration of the cells.^39-41^ Chemokine CXCL 7 is engaged in neutrophil-platelet crosstalk, and also it is actively involved in the growth of renal cell carcinoma.^42^ IL10 Rb is the receptor for IL10, and it actively participates in inflammatory signaling.^43, 44^ IL 10 regulates memory T cell development in response to viral infections.^45^ Eotaxin 2 is an eosinophilic chemoattractant protein, and it acts through CCR3, and it is actively involved in the recruitment of other inflammatory cells also.^46^ BMP 7 acts on the BMP receptor, which is actively involved in the process of inflammation and atherogenesis. Exogenouos administration of BMP7 improves left ventricular remodeling and function in acute myocardial infarction.^47^

Trappin 2 is a serine protease inhibitor, and it has anti-inflammatory actions on the mucosal surfaces.^48^ It also has anti-retrovirus activities on the mucosal surfaces. SDF1 alpha and its chemokine receptor play a significant role in hematopoietic cell mobilization, cancer metastasis, and ischemic injury repair in myocardial infarction tissues.^49^ FGF 7 (fibroblastic growth factor) has an active role in tissue repair.^50^ CCL 28 is a mucosa-associated epithelial chemokine, and it is associated with the recruitment of the cells, and it helps in T and B cell accumulation in mucosal surfaces.^51^ 4-1BB (CD137) signalosome promotes T cell proliferation and survival and results in increased T cell effector functions.^52,53^ VCAM1 is an inflammatory protein involved in cell to cell adhesions, and it also effectively induces angiogenesis.^54,55^

GITR Glucosteroid induced TNFR related protein is expressed by T cells and its ligands, and it boosts T cell activity. GITR agonistic stimulation is emerging as a promising therapeutic concept.^56^ SIGLEC 5 is a leucocyte receptor that recognizes sialic acid structures and helps in leucocyte recruitment.^57^ VEGFR3 is a receptor for vascular endothelial growth factors C and D and it is involved in lymphangiogenesis and to some extent in VEGF A induced angiogenesis as well.^58^ IL13 overcomes insulin resistance by promoting anti-inflammatory macrophage differentiation in adipose tissue.^59^ CD30 is expressed on the surfaces of the endothelial cells though it is primarily expressed by lymphoid tissues. They are expressed in non lymphamatous tumours. CD30 signalling is involved in proliferation, differentiation and survival (anti-apoptosis).^60^ TGF B2 is expressed in the endothelium, and it plays an essential role in angiogenesis.^61^ GP130 is a glycoprotein that participates in IL6 mediated inflammation and vascular pathologies, and it also has a negative feedback control.^62^

### Reduced biomarkers with lysate

MIF is a widely expressed pleiotropic cytokine, and it is involved in the stimulation of other inflammatory cytokines like TNF alpha, INF gamma, IL6, IL 12, CXCL8 etc.^23,63^ MIF is elevated in type 1 and 2 diabetes. Fas activation is associated with autoimmune disorders, which can be modulated by downregulation.^64^ TIMP 2 are matrix metalloproteinases which are involved in inflammation in the cancer cells.^65^ IGFBP2 are inhibitory and stimulatory to some of the tumours.^66^ IL13 R2 is involved in mediating inflammation leading to myocarditis. Hence a reduction in these receptors could reduce inflammatory changes.^67,68^ Dickkopf 1 family proteins are active modulators of Wnt pathways, and mostly their effects are inhibitory.^69^ TNF R2 has proinflammatory and some anti-inflammatory aspects as well. The stimulatory and inhibitory effects had attracted considerable interest in the treatment of autoimmune diseases and cancer.^70^ Follistatin is actively involved in activin A-follistatin regulation of cardiac inflammation and fibrosis.^71^ Heparin-Binding Epidermal Growth Factor-Like Growth Factor Inhibits Cytokine-Induced NF-κB Activation.^72^ The heat map analysis shows a significant change in more proteins, and thereby it is possible to infer that streptococcus has a role in immune regulation.

The above observations indicate that the immune system undergoes various modifications by the streptococcus pyogenes direct challenge. Certain parameters are increased, and some are decreased. In the long term, the immune memory and its regulation are complex, and it is also subjected to many positive and negative feedback regulations. Also, when the lysate is added, the modulations are seen. Hence, streptococcus pyogenes is associated with changes in the immune system, which can influence potential regulations in the immune homeostasis of the individuals. The negative impact of rheumatic heart diseases can have a positive influence in modifying the immune-related functions and possibly rendering significant protection.

### Chaos and decoding

It is indeed difficult to predict the immune response in the future to a certain extent. However, it can be inferred that the endothelial response could be significant and probably chaotic, which determine transcription and gene regulation.^73^ Chaos does not necessarily lead to dysfunctions, and at times and situations, it could be a natural method of selection to strengthen immune functions. Optical laser chaos signals which are high speed when studied are found to synchronize and it has many features.^74, 75^ Decoding these chaotic signals in immune system would potentially lead to our better understanding of immune system and modify its treatment which are significant challenges to at present.

### Streptococcus vs. viral infections – both together forever

In the recent times coronavirus infections (Covid-19) pandemic is rampant, and the infection selectively affects various countries, and the mortality statistics were varied in different countries. These could be due to varied climatic conditions and the immune response to the viruses. The Southeast Asian countries and the Indian subcontinent are relatively less affected so far, at his time of writing. The streptococcus pyogenes, tropical bacterial infections and other viral are common in these areas, and they could provide cross-immunity to the coronaviral infections.^76^ Bacteria can synthesize restriction enzymes like nucleases and inhibit viruses.^77^ It has been shown that bacterial presence can reduce the intensity of viral infections.^78^ Our study also reflects the immune regulation changes due to streptococcus pyogenes as well as by its lysate.

### Hygiene and the Dirt

In the theory of natural selection, the environment or the Nature in various forms could offer protection by its selective mechanisms in various geographic locations.^79^ These could not necessarily be simple mechanisms but also as chaotic or cross-immunity methods. The commonly available streptococcus bacteria and the common viruses could be the mechanism of choice in the form of selection by unhygienic means.^80^ Possibly these protective mechanisms are not well studied by the scientific and social communities, and the primary focus is often tertiary care treatment modalities.

### Metabolic protection and autoimmunity regulation

The rarity of diabetes and rheumatic heart disease was observed by legendary physicians like Joslin and Steinchron in the early 1920s and 1930’s respectively.^81,82^ Our study also suggests various metabolic modulators being stimulated and some inhibited. Also, Joseph Barach made similar observations in that period and attributed the views to immune regulation changes. Our observations, such as increased IL13, decreasing Fas, and Dickkopf proteins, also indicate possible metabolic protection. Antibodies seen in rheumatic fever are also found in antiphospholipid antibody syndromes.^83^ However, the incidence of autoimmune diseases in rheumatic heart disorders is very rare or possibly mutually exclusive by negative feedback mechanisms, at least in the clinical experience of the author.

### Limitations and future perspectives

Further studies need to be performed to observe the immune changes in animals after direct streptococcus challenge and after the lysate administration. Specifically, the immune changes regulating the autoimmune disorders, cancer regulation, atherosclerotic processes, and host defense activities to viruses need to be studied in animal models.

## Conclusion

Streptococcus pyogenes and its lysate has immunomodulation actions when tested with endothelial cells, which have pleiotropic functions. Further studies need to be performed to identify its potential benefits.

## Ethical considerations

Not applicable

## Author contributions

MCA conceived the idea and method, designed the study, interpreted results and wrote the paper. JW prepared the methods protocol, performed the tests, derived the results and principal component analysis.

## Footnotes

The study was presented as an abstract in Synthetic Biology UK 2019.

**Tables 1 to 5 shows biomarkers expression after direct streptococcus pyogenes injury on the endothelial cells.**

